# Neural Readout of a Latency Code in the Active Electrosensory System

**DOI:** 10.1101/2021.12.14.472594

**Authors:** Krista E. Perks, Nathaniel B. Sawtell

## Abstract

The latency of spikes relative to a stimulus conveys sensory information across modalities. However, in most cases it remains unclear whether and how such latency codes are utilized by postsynaptic neurons. In the active electrosensory system of mormyrid fish, a latency code for stimulus amplitude in electroreceptor afferent nerve fibers (EAs) is hypothesized to be read out by a central reference provided by motor corollary discharge (CD). Here we demonstrate that CD enhances sensory responses in postsynaptic granular cells of the electrosensory lobe, but is not required for reading out EA input. Instead, diverse latency and spike count tuning across the EA population gives rise to graded information about stimulus amplitude that can be read out by standard integration of converging excitatory synaptic inputs. Inhibitory control over the temporal window of integration renders two granular cell subclasses differentially sensitive to information derived from relative spike latency versus spike count.

## INTRODUCTION

The latency of spikes evoked by sensory stimuli convey information about non-temporal features (e.g. stimulus amplitude, location, or identity) at various processing stages across sensory modalities, including in vision (Gawne et al., 1996; Gollisch and Meister, 2008; VanRullen et al., 2005), audition (Ashida and Carr, 2011; Chase and Young, 2007; Furukawa and Middlebrooks, 2002; Grothe and Klump, 2000; Heil, 2004; Zohar et al., 2011), olfaction (Bathellier et al., 2008; Cury and Uchida, 2010; Shusterman et al., 2011), somatosensation (Johansson and Birznieks, 2004; Panzeri et al., 2001; Saal et al., 2016), and active electrosensation (Bell, 1990b; Hall et al., 1995; Szabo and Hagiwara, 1967). Latency codes have potential advantages over conventional rate codes in terms of speed (Gollisch and Meister, 2008; VanRullen et al., 2005), information capacity (Rieke et al., 1996; Theunissen and Miller, 1995), and energy efficiency (Lennie, 2003). Moreover, latency information appears to be sufficient for aspects of olfactory (Chong et al., 2020; Chong and Rinberg, 2018; Smear et al., 2011), tactile (Thomson and Kristan, 2006), and electrosensory (Hall et al., 1995) mediated behavior. However, unlike conventional spike rate codes, latency codes ostensibly require explicit postsynaptic readout mechanisms, the nature of which remains controversial (Stanley, 2013). Motor corollary discharge (CD) has been hypothesized to provide a central reference signal for reading out latency codes in cases in which sensory input is time-locked to behavior (Bell, 1989; Crapse and Sommer, 2008; Moore et al., 2013; Shusterman et al., 2011). Alternatively, latency codes could be read out based on information contained in the relative timing of spikes across a population of inputs (Haddad et al., 2013; Panzeri et al., 2014; Uchida et al., 2014; Zohar and Shamir, 2016).

The active electrosensory system of weakly electric mormyrid fish offers a number of advantages for examining how latency information is utilized by postsynaptic neurons. Mormyrid fish emit brief pulsed electrical fields known as electric organ electric discharges (EODs). Nearby objects with conductivity higher (lower) than the surrounding water increase (decrease) the amplitude of the EOD-induced field on the skin (Figure 1A). A-type mormyromast electroreceptors on the skin transduce increases (decreases) in the local amplitude of the EOD pulse into highly-precise, smoothly graded decreases (increases) in spike latency (Bell, 1990b; Sawtell et al., 2006; Szabo and Hagiwara, 1967). Although most EAs also fire more spikes in response to EOD amplitude increases, evidence from intra-axonal recordings (Bell, 1990a), information theoretic analysis of EA responses (Bell, 1990b; Sawtell and Williams, 2008), and behavioral studies (Hall et al., 1995) suggest the functional importance of latency coding in this system. EAs project somatotopically to the hindbrain electrosensory lobe (ELL) where they form excitatory synapses predominantly with two anatomically distinct subclasses of interneurons known as deep and superficial granular cells (DGCs and SGCs) (Bell et al., 2005; Bell et al., 1989; Zhang et al., 2007) (Figure 1B). Granular cells send their axons to the superficial layers of the ELL where they synapse onto the output cells of the ELL and hence serve as an obligatory relay for electrosensory information (Bell et al., 2005). CD inputs related to the EOD motor command are prominent in the ELL and have been hypothesized to provide a reference signal that could serve to read out latency-coded input at the level of the ELL granular cells (Bell, 1989, 1990a; Hall et al., 1995) (Figure 1C). However, the small size and dense packing of granular cells has, until now, prevented *in vivo* recordings required to test this hypothesis.

**Figure 1.**
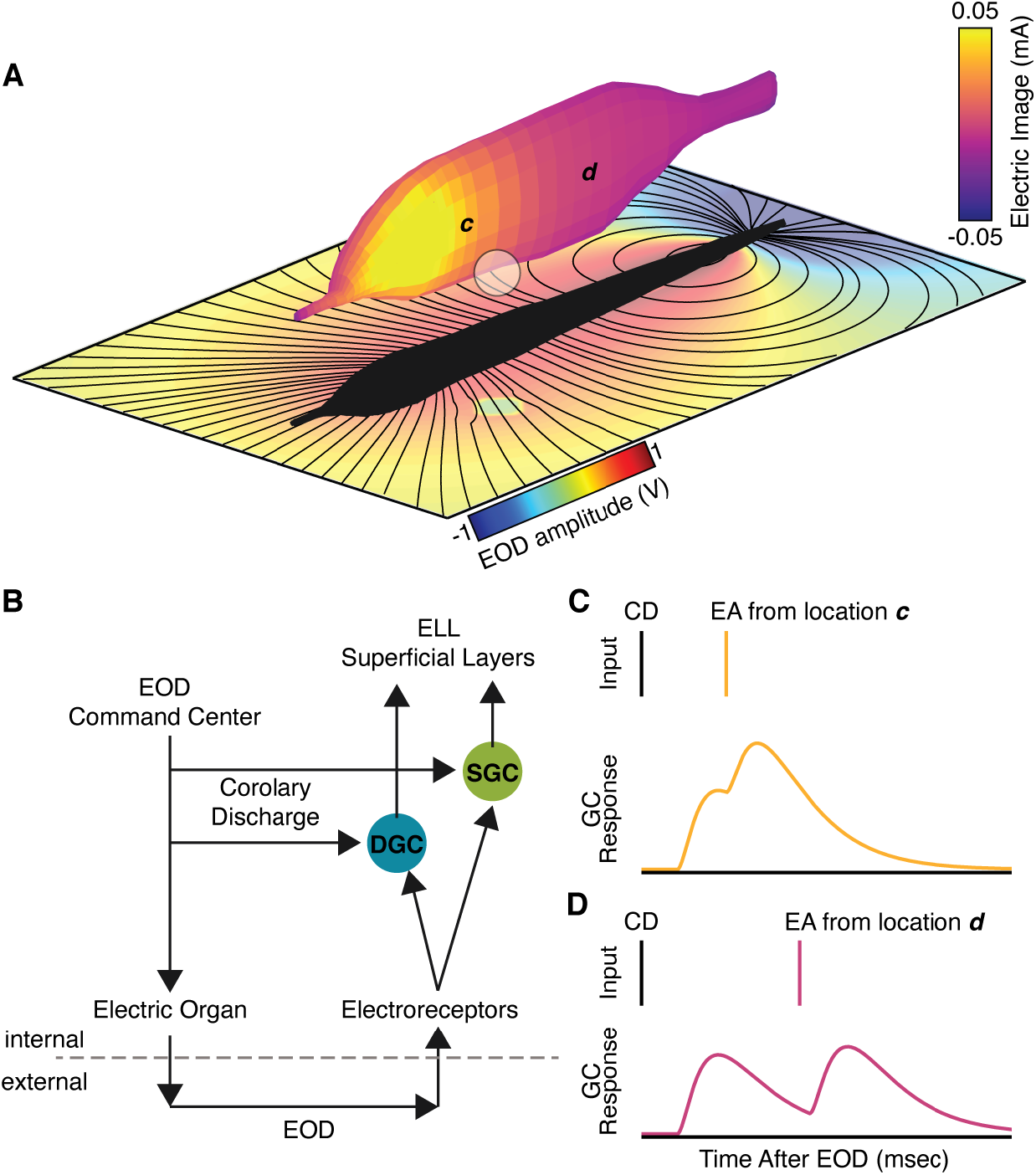
Central readout of a latency code for electrosensory stimulus amplitude based on corollary discharge. (A) Weakly electric mormyrid fish emit brief pulses of electricity known as electric organ discharges (EODs; bottom plane). Nearby objects induce “electrical images” on the body surface by changing local current density (amplitude) of the resulting electric field. Electrical image amplitude is encoded by the latency and number of spikes fired by afferent fibers innervating electroreceptors on the skin (EAs). (B) Two distinct classes of granular cells in the electrosensory lobe - superficial (SGC; green) and deep (DGC; teal) - integrate excitatory input from EAs with a centrally-originating corollary discharge (CD) input related to the motor command to discharge the electric organ. (C) Latency coded information is hypothesized be read out in granular cells (GCs) based on summation of EA inputs (yellow and purple lines) with a fixed temporal reference signal provided by CD (black lines). In such a scheme, a short latency EA spike arriving near the peak of the CD-evoked depolarization (yellow) would yield a larger amplitude granular cell postsynaptic response than a longer latency spike arriving on its falling phase (purple).

## RESULTS

### Two physiologically distinct granular cell subclasses integrate EA and CD input

Granular cell responses were characterized using blind whole-cell recordings from the medial zone of the ELL in awake paralyzed fish (STAR Methods). In this preparation, neuromuscular paralysis blocks the EOD, but EOD motor commands continue to be emitted spontaneously by the fish at a rate of ~3-5 Hz, leaving CD input to the ELL intact. The EOD is mimicked by a brief electrical pulse delivered either at the naturally occurring 4.5 ms delay relative to the EOD motor command or at a long (50 ms) delay. The former condition (termed short delay) is used to study the normally occurring interactions between electrosensory and CD inputs while the latter (termed long delay) allows sensory and CD inputs to be observed separately.

Histological recovery of a subset of recorded granular cells filled with biocytin revealed laminar locations and morphological properties consistent with previous anatomical descriptions of DGCs and SGCs (Bell et al., 2005; Zhang et al., 2007) (Figure 2A). We report exclusively on subthreshold responses because somatically recorded action potentials were small and often difficult to distinguish, likely due to an electrotonically remote site of initiation at the distal end of a thin initial segment (Bell et al., 2005; Zhang et al., 2007). Electrosensory stimulation evoked short-latency synaptic excitation in both DGCs and SGCs, consistent with monosynaptic input from EAs. Excitation in DGCs was sharply peaked and appeared to be truncated by inhibition (Figure 2B,D, *blue arrow*), while excitation in SGCs decayed much more slowly (Figure 2C,D). Prior electron microscopy and *in vitro* studies indicate that GABAergic interneurons, known as large multipolar intermediate layer (LMI) cells, form inhibitory synapses onto granular cells (Han et al., 2000; Meek et al., 2001; Zhang et al., 2007). Differences in the time course of electrosensory responses in DGCs and SGCs may reflect differences in the strength of LMI-mediated inhibition.

**Figure 2.**
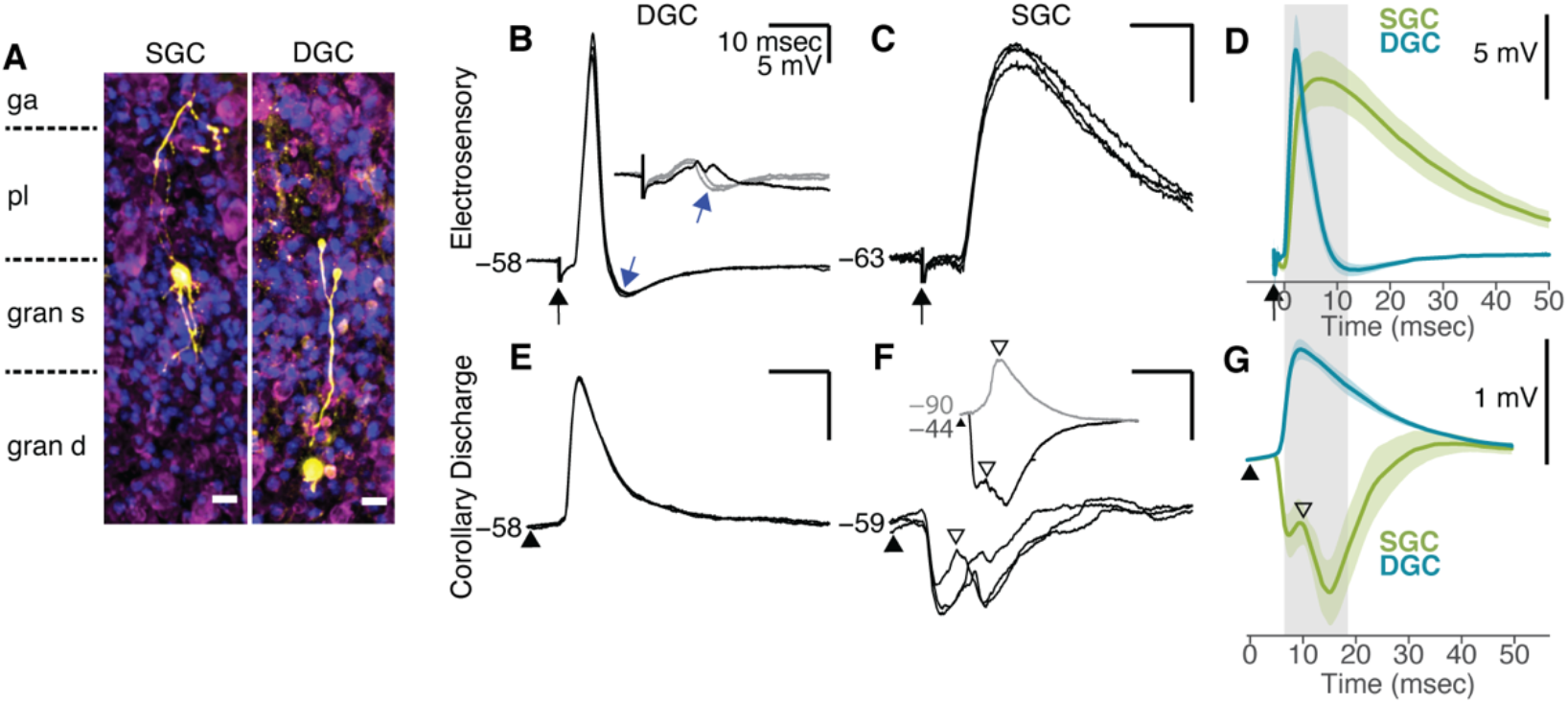
Responses of granular cells to electrosensory and corollary discharge inputs. (A) Confocal z-stacks of an SGC and a DGC labeled with biocytin (yellow) during whole cell recording. DAPI (blue) and Neurotrace (magenta) staining show the layers of the ELL. Scale bar, 10 μm (gang, ganglion layer; plex, plexiform layer; gran s, superficial granular cell layer; gran d, deep granular cell layer). (B) Example DGC response to electrosensory input (three trials overlaid) aligned to stimulus onset (black arrow). Baseline membrane potential was −58 mV. The rapid decay of the EPSP is likely due to synaptic inhibition (blue arrow). Inset: three trials overlaid in response to a weaker electrosensory stimulus at threshold for evoking an IPSP (blue arrow) that truncates the electrosensory-evoked EPSP. The black trace shows a trial in which the EPSP occurred in the absence of the IPSP. (C) Example SGC response to electrosensory input (three trials overlaid) aligned to stimulus onset (black arrow). Baseline membrane potential was −63 mV. (D) Average response of DGCs (teal, n=24) and SGCs (green, n=18) to the electrosensory stimulus. Responses aligned to the EPSP onset (time=0) to average across cells. Shading denotes SEM. DGC responses (n = 24) peaked earlier (2.5 ± 1 ms versus 11 ± 8.2 ms; t(42)=−4.99, p< 0.001), decayed more rapidly (2.6 ± 1.3 ms versus 33.3 ± 32.8 ms; t(42)=−4.61, p< 0.001), and were narrower (half-width at half-height 3.6 ± 1.7 ms versus 24.8 ± 11.4 ms; t(42)=−9.04, p< 0.001) than those of SGCs (n = 18). (E) Example DGC response to electric organ corollary discharge (CD) input (three repeated trials overlaid). Triangle indicates the time of the EOD motor command recorded by an electrode near the electric organ. (F) Example SGC response to CD input (three repeated trials overlaid). Inset: CD response for a different SGC illustrating an excitatory response evident at a hyperpolarized membrane potential near reversal for the inhibitory response in this cell (gray; trial average). At a depolarized membrane potential, this cell showed rapid, early onset inhibition (black; trial average). (G) Average CD responses across a subset of DGCs (teal; n = 5) and SGCs (green; n = 5) selected for their similar resting membrane potentials (−50 to −60 mV). Shading denotes SEM. Gray boxed region indicates the range of latency shifts observed in EAs (6.5 to 18.5 ms), shown relative to the timing of granular cell CD responses. CD input evoked short-latency EPSPs in DGCs (onset 5.7 ± 0.8 ms; peak 9.7 ± 2.7 ms, n=24) and even shorter latency IPSPs in SGCs (4.9 ± 0.5 ms, n=12). A depolarizing PSP was observed after inhibition onset (open triangle; onset 7.6 ± 2.2 ms; peak 14.4 ± 5.8 ms) in 12/18 SGCs. In 3/24 DGCs we observed a prominent IPSP at around 7.7 ms.

The possibility that granular cells receive CD input related to the EOD motor command has been suggested based on prior work but not directly shown (Bell, 1990a). Our intracellular recordings confirmed prominent CD input to granular cells. DGCs exhibited highly-stereotyped, short-latency excitation time-locked to the EOD motor command (Figure 2E,G, Figure S1). In some DGCs, CD excitation appeared to be truncated by inhibition (see Figure 3C, black trace), similar to their responses to electrosensory input. In contrast, SGCs typically exhibited a stereotyped, short-latency hyperpolarization followed by a depolarization (Figure 2F,G, Figure S1). At more hyperpolarized potentials, CD responses consisted mainly of a depolarization (Figure 2F, *inset, open arrowheads*), suggesting that CD input to SGCs is comprised of synaptic inhibition followed by excitation. Prior studies suggest that CD input to the ELL originates from the medial juxtalobar nucleus (JLm) (Bell and von der Emde, 1995; Mohr et al., 2003). Consistent with this, electrical microstimulation in the vicinity of the JLm evoked synaptic responses resembling naturally-occurring CD responses, including short-latency depolarizations in DGCs and hyperpolarizations in SGCs (Figure S1). However, since JLm neurons are thought to be glutamatergic (Bell and von der Emde, 1995; Mohr et al., 2003), the origin of CD-inhibition in SGCs is unclear.

**Figure 3.**
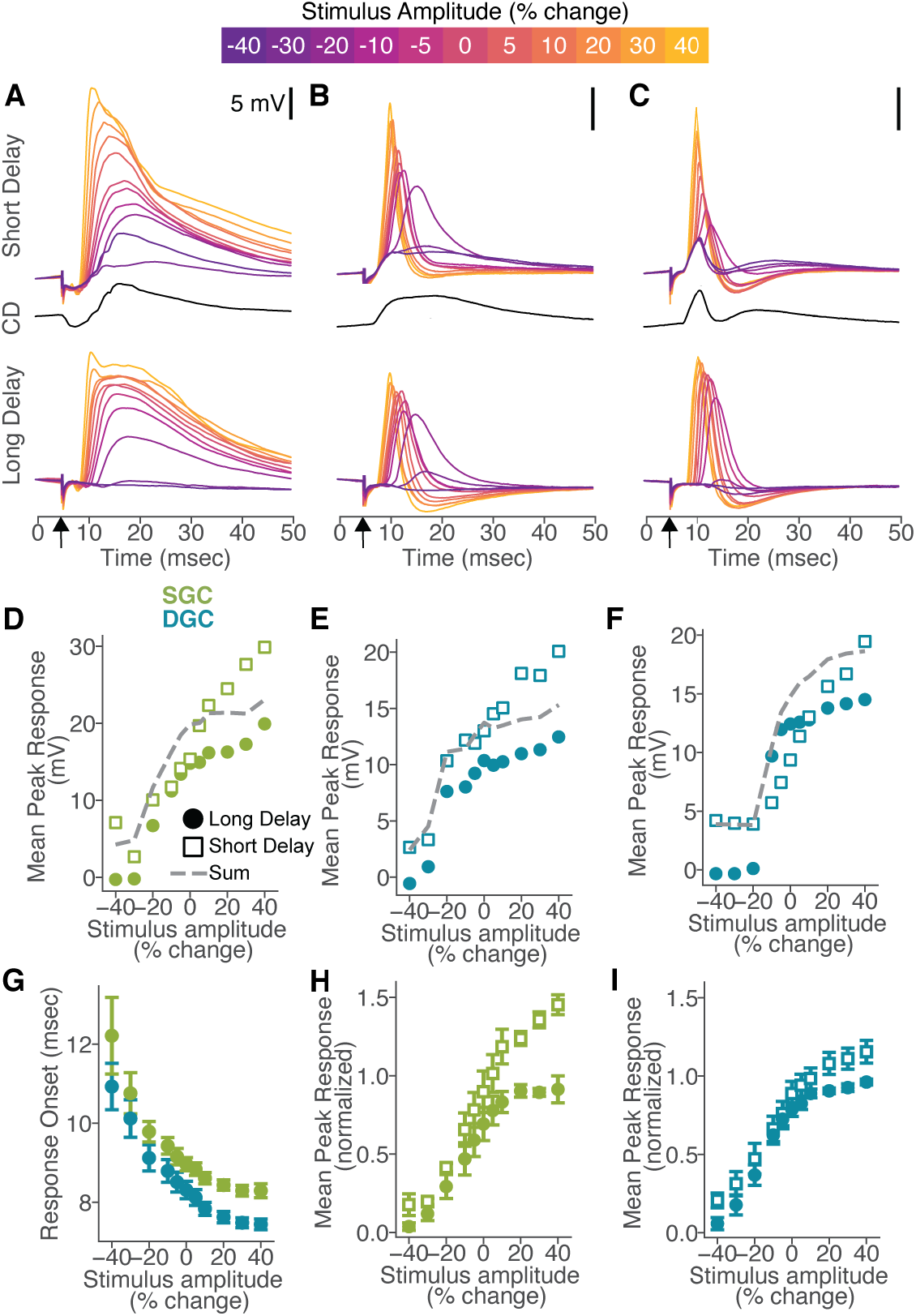
Corollary discharge enhances but is not required for electrosensory responses in granular cells. (A) Responses of an example SGC across amplitudes (color bar, top) to stimuli delivered either at a short (top) or long (bottom) delay relative to the EOD motor command. Middle: average response to corollary discharge (CD) input alone. Arrow indicates time of the electrosensory stimulus. (B-C) Same displays as in A, but for two example DGCs. (D) Mean peak response versus stimulus amplitude for the same SGC shown in A in long delay (filled circle) versus short delay (open square) conditions. The predicted response at a short delay (‘sum’, gray dashed) was calculated by adding the long delay response at each stimulus amplitude to the CD response. (E,F) Same displays as in D but for the example DGCs shown in B and C. (G) Response onset latency (relative to stimulus onset; mean ± SEM) versus stimulus amplitude for SGCs (green; n=5) and DGCs (teal; n=13). (H) Peak response amplitude versus stimulus amplitude under long delay (filled circle) versus short delay (open square) conditions for SGCs (n = 5; mean ± SEM). Responses within each cell were normalized by the maximum long delay response amplitude before averaging across cells. The delay between the electrosensory stimulus and the command had a significant effect on peak response amplitude, F(1,88) = 55, *P* < 0.001 (two-factor repeated measures ANOVA). Across the range of −10 to 10% stimulus amplitude, the sensitivity at a long delay was 8.3 ± 5.6 mV and the sensitivity increased to 27.9 ± 8.8 mV at a short delay. Out of 4 SGCs with at least 5 trials in every condition, all had a significant effect of stimulus delay on raw peak response amplitude (p< 0.001). (I) Same as in H, but for DGCs (n = 13; mean ± SEM). The delay between the electrosensory stimulus and the command had a significant effect on peak response amplitude, F(1, 264) = 21, *P* < 0.001 (two-factor repeated measures ANOVA). Across the range of −10 to 10% stimulus amplitude, the sensitivity at a long delay was 8.1 ± 8.1 mV and the sensitivity increased to 26.8 ± 14.6 mV at a short delay. Out of 9 DGCs with at least 5 trials in every condition, 7 had a significant effect of stimulus delay on raw peak response amplitude (p< =0.001). (H) and (I): Significant effects of stimulus amplitude and delay, but not delay-by-amplitude interaction at P < 0.001 on the mean peak response amplitudes across cells.

### CD input enhances but is not required for reading out EA input to granular cells

Next we characterized granular cell responses to modulations of electrosensory stimulus amplitude. Such modulations mimic increases and decreases in EOD amplitude due to conducting and nonconducting objects, respectively (STAR Methods). To test the role of CD input, we compared granular cell responses evoked by electrosensory stimuli delivered at short (naturally-occurring) versus long delays relative to the EOD command. In both conditions, increases in electrosensory stimulus amplitude led to decreases in postsynaptic response onset in granular cells (Figure 3A-C,G), as expected based on well-characterized latency shifts in EAs (Bell, 1990b; Szabo and Hagiwara, 1967). Large graded changes in postsynaptic response amplitude (>10 mV) were observed in response to stimuli presented at both short and long delays (Figure 3A-C,H,I). These findings are consistent with the hypothesis that latency coded EA input induces changes in postsynaptic response amplitude in granular cells but suggest that CD input is not required for this transformation.

To further evaluate the effect of CD input, we compared measured granular cell responses to those calculated based on a linear sum of the granular response to electrosensory stimuli delivered at a long delay and the response to the CD input alone (Figure 3D-F, *dashed line*). Most SGCs exhibited supralinear summation of electrosensory and CD input, particularly at high stimulus amplitudes (Figure 3A,D,H; Figure S2). The enhancement of sensory-evoked depolarizations in SGCs is notable given that SGCs typically exhibited CD-evoked inhibition (Figure 3A; Figure S2). Supralinear summation was not observed between CD input and EPSP waveforms generated by somatic current injections in SGCs (Figure S2), suggesting that facilitatory interactions between CD and EA input might occur electrotonically distant from the soma in the long, thin dendrites of SGCs (Bell et al., 2005; Zhang et al., 2007). More varied interactions between CD and electrosensory inputs, including both supralinear (Figure 3B,E) and sublinear (Figure 3C,F) summation, were observed in DGCs. Sublinear summation is expected for cells, like the example shown in Figure 3C, in which CD excitation is followed by inhibition. On average, DGC responses were modestly enhanced by CD input (Figure 3I).

Finally, we recorded subthreshold responses to modulations of electrosensory stimulus amplitude in E-type output cells of the ELL, one of the major postsynaptic targets of granular cells (Bell et al., 2005; Grant et al., 1996). Responses of E cells resembled those of granular cells in that long delay electrosensory stimuli evoked graded changes in subthreshold response magnitude that were enhanced when stimuli were delivered at the natural delay relative to the EOD motor the command (Figure S3). Overall, these results argue against the hypothesis that CD is required for reading out electrosensory input, but are consistent with a role for CD in enhancing or “gating-in” responses to the fish’s own EOD (Meyer and Bell, 1983).

### Diverse spike latency and number tuning in EAs

Granular cells pool excitatory input from an estimated 4-7 EAs innervating nearby electroreceptors on the skin (Bacelo et al., 2008; Bell, 1990a). To determine the potential implications of such convergence for reading out electrosensory input, we obtained intracellular recordings from EAs in the granular layers of the ELL near their central terminals. Responses to identical electrosensory stimuli were obtained from 7-21 individual EAs in each of four fish. Recording locations were restricted to the same somatotopic region of the ELL to minimize potential response variability due to electroreceptor location on the skin. All EAs exhibited smoothly graded decreases in spike latency with increases in stimulus amplitude and most also exhibited increases in spike number (Figure 4A-C), consistent with prior studies (Bell, 1990b; Sawtell et al., 2006; Szabo and Hagiwara, 1967). Although responses were highly reliable across repeated trials within individual EAs, substantial heterogeneity was observed in latency tuning across the population (Figure 4A-C). To quantify this, we fit exponential functions to first spike latency-stimulus amplitude response curves for all recorded EAs (Figure 4D) and plotted the distribution of parameters for each fish (Figure 4E-G). Substantial heterogeneity was observed across EAs within each fish in terms of the sensitivity of latency shifts to stimulus amplitude (decay), the total range of latency shifts (amplitude), and the minimum first spike latency (offset). To isolate effects of relative latency shifts from recruitment, we identified the range of stimulus amplitudes over which EAs fire at least one spike (Figure 4H, gray box). Because first spike latencies converge onto a minimum value, presumably set by biophysical limits, heterogeneous latency tuning results in a stimulus-dependent decrease in the interval between first spikes across the population (Figure 4H). This implies that a rate code for stimulus amplitude exists at the level of the input population even when the total number of EA spikes remains constant.

**Figure 4.**
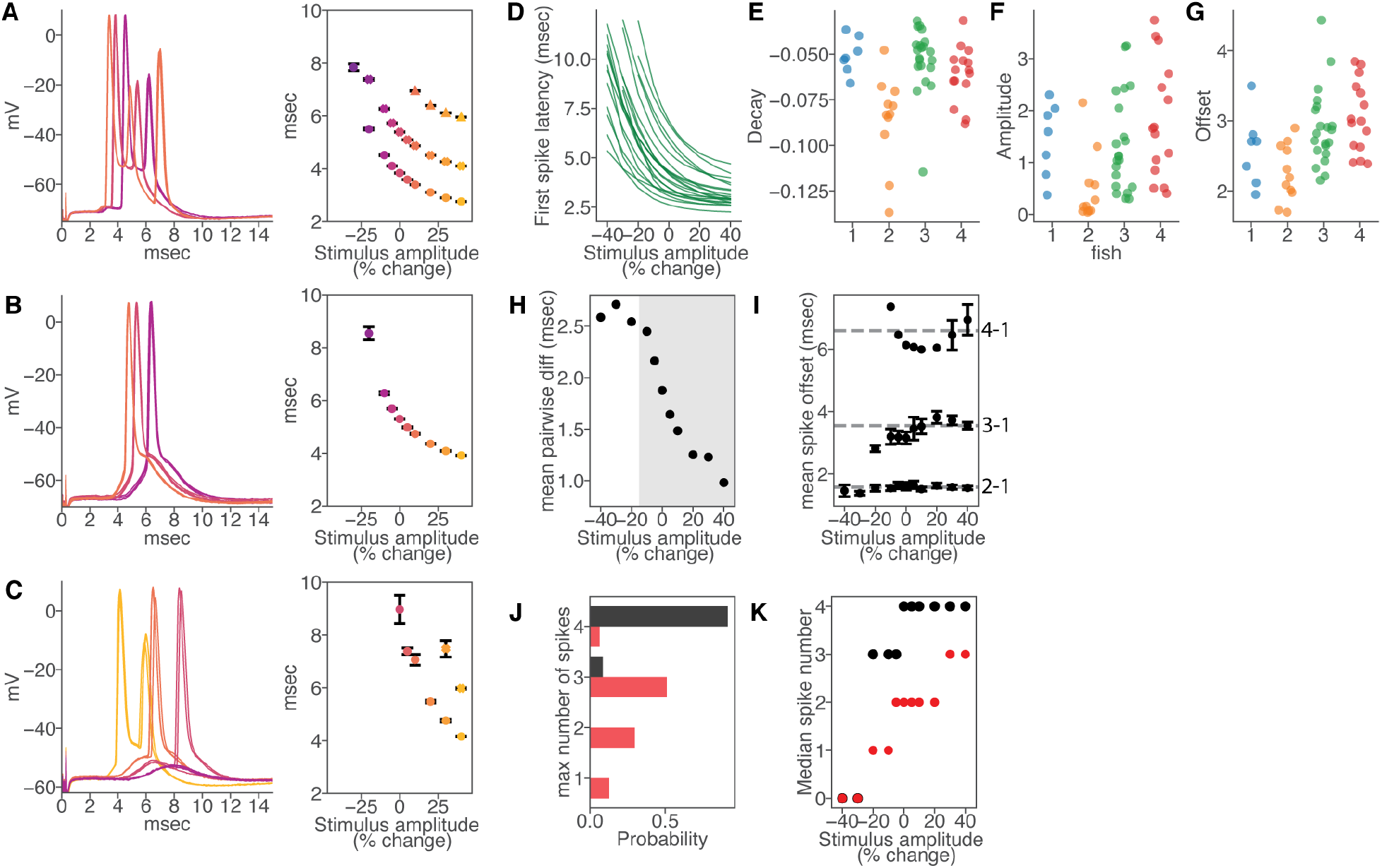
Electroreceptor afferents exhibit diverse first spike latency and spike count tuning. (A) Left: Intracellular responses of a recorded EA at stimulus amplitudes of −10, 0, and +10 % (colored according to the scatterplot at right). Three trials overlaid at each amplitude. Right: First spike latency versus stimulus amplitude (mean ± SEM). (B,C) Same as A, but for two other example EAs with notable differences in latency and spike count tuning. In (C), three trials at +40 % amplitude are also shown. (D) Exponential fits of first spike latencies across stimulus amplitudes for 21 electroreceptor afferents recorded from a single fish (fish 3 in E-G). (E-G) Heterogeneity of fit parameters among EAs recorded in four fish (n=7,11,21,15 EAs per fish). (H) Mean pairwise latency among first spikes (n=58 total EAs in 4 fish). Shading denotes stimulus range for which >94% of EAs have at least one spike. (I) Scatterplot of spike offset (mean ± SEM) for second (1.6 ± 0.5 msec, n= 360 spikes), third (3.6 ± 0.8 msec, n= 145 spikes), and fourth (6.6 ± 0.8 msec, n= 13 spikes) spikes relative to first spike latency. For each datapoint (mean ± SEM), EAs without that spike number were ignored. (J) Probability of EAs having 0, 1, 2, 3, or 4 spikes at the maximum stimulus amplitude (+40%) for recorded data (red) and the prediction (black) based on recorded first spike latency and mean spike offsets (as in I) (n=58 EAs). (K) Median number of spikes per EAs across stimulus amplitude for actual (red) and predicted (black) spikes (n=58 EAs).

EAs also exhibited diversity in their spike count tuning (Figure 4A-C). Consistent with prior results, second and subsequent spikes followed earlier spikes by a fixed offset (Figure 4I) (Bell, 1990b; Sawtell et al., 2006). Notably, the number of spikes fired by an individual EA appeared unrelated to its first spike latency tuning. To demonstrate this, we compared the actual distribution of maximum spikes fired by each EA (Figure 4J, *red*) to a simulated distribution in which subsequent spikes were simply added at a fixed delay after prior spikes up to a maximum latency estimated from the data (Figure 4J, *black*). Recorded EAs exhibited a broader distribution of total spikes than expected based on the simulation, suggesting that spike latency and spike count tuning vary independently within EAs. Comparing the median EA spike count as a function of stimulus amplitude for recorded versus simulated populations, suggests that this independent tuning results in spike count grading over a wider range of stimulus amplitudes (Figure 4K).

### Postsynaptic readouts in SGCs and DGCs based on EA convergence

The foregoing results suggest that the EA population contains graded information about stimulus amplitude that could be read out based on simple input summation in granular cells independent of a fixed reference signal. To test this, we constructed conductance-based model neurons with parameters adjusted to match the rise and decay of near-threshold EPSPs recorded in SGCs and DGCs (Figure S4). Input to the model consisted of spikes from 4 EAs randomly subsampled from the recorded population. Though we chose to focus on simplified models to gain insight into the functional significance of EA convergence, numerous additional factors may contribute to measured subthreshold responses in granular cells, e.g. mixed chemical and electrical synapses between EAs and granular cells and voltage-gated conductances (Zhang et al., 2007). Responses in model cells receiving only excitatory EA input were broad in duration and exhibited large, graded increases in peak depolarization as a function stimulus amplitude, similar to recorded SGCs (Figure 5A,B). To mimic the brief electrosensory responses exhibited by DGCs, we added strong inhibitory input at a short delay (2-4 ms) after the onset of the excitatory response, effectively truncating the time window for integrating EA input. Model responses were much briefer under these conditions, resembling recorded DGCs, but still exhibited large, graded increases in peak depolarization as a function stimulus amplitude (Figure 5C,D). Aside from changes in the absolute magnitude of responses, results were similar over a range of values for EA input number and inhibitory input delay (Figure S4). These results suggest that granular cell responses to electrosensory stimuli observed *in vivo* can largely be explained based on convergence of a small numbers of excitatory EA inputs and, for DGCs, sensory-evoked inhibition.

**Figure 5.**
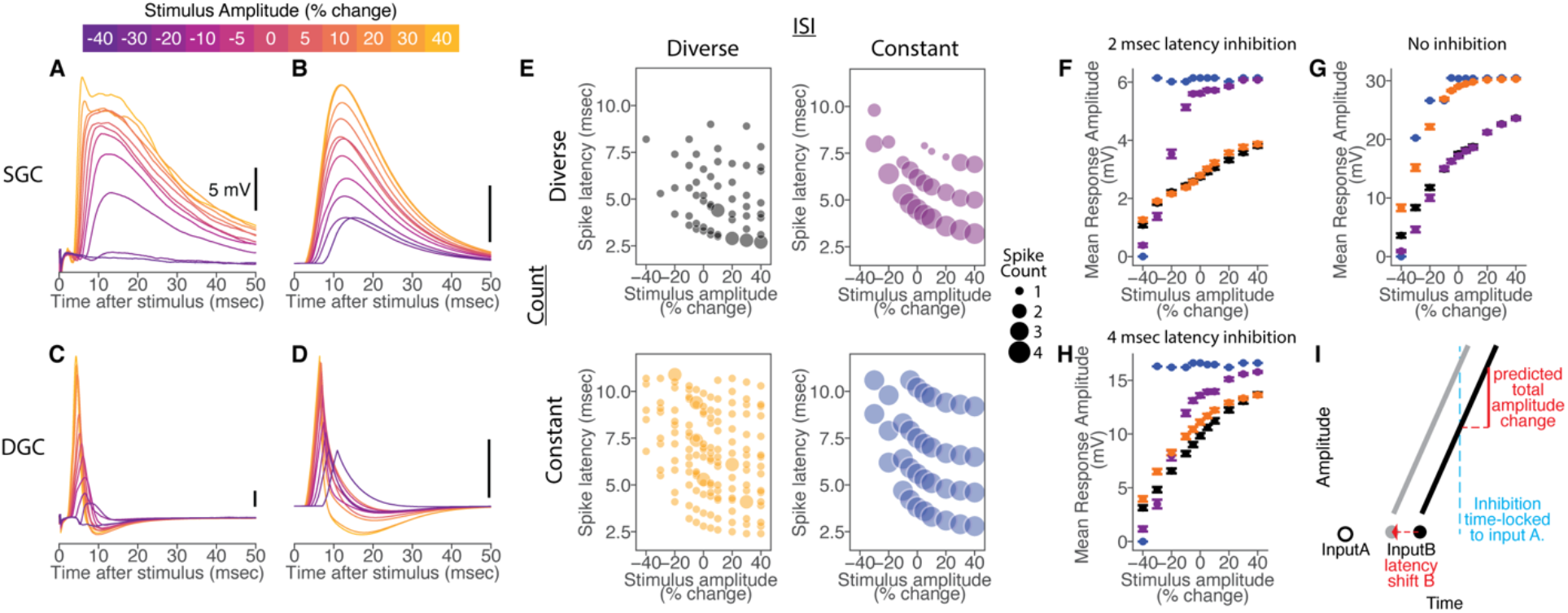
Summation of diversely tuned EAs accounts for graded readout of stimulus amplitude in granular cells. (A) Responses of an example SGC to electrosensory stimuli of different amplitudes (color bar, top) delivered at a long delay from CD input. (B) Example responses of a model granular cell receiving 4 excitatory EA inputs. (C) Responses of an example DGC to electrosensory stimuli of different amplitudes delivered at a long delay from CD input. (D) Example responses of a model granular cell with the same excitatory EA input as in B, but with an inhibitory conductance driven by the first spike in the EA input population, at a 4 ms delay. (E) Examples of four sets of EA population inputs (n = 4 EAs per population) designed to test the effects of diverse latency and spike count tuning on granular cell responses (see main text). Circles denote the time of EA spikes with the size indicating the number of spikes. Black, subsampled recorded EA data; Orange, diverse latency tuning is preserved while diverse spike count tuning is removed; Purple, diverse latency tuning is removed while diverse spike count tuning is preserved; Blue, diverse latency and spike count tuning are both removed from the input population. (F) Peak response amplitude versus stimulus amplitude across model DGCs (mean ± SEM; n = 200 different EA input populations) under each condition shown in E with inhibition delayed by 2 ms relative to the first EA spike. (G) Same as in F, but for model SGCs. (H) Same as in F, but with inhibition delayed by 4 ms relative to the first EA spike. (I) Shifts in a second EA input *B* relative to a first EA input *A* (dotted red arrow) cause a predictable change in the amplitude of the second EPSP when an inhibitory input (blue), time-locked to input *A*, cuts off the response during the EPSP rise. The predicted response amplitude (red, solid line) change depends on the slope of the EPSP rise.

Next, we used the model to test the respective contributions of diversity in latency versus spike count tuning in EAs to postsynaptic responses in granular cells. Model SGC and DGC responses to subsampled EA inputs were compared for three sets of manipulated EA inputs in which: (1) diversity in both latency and spike count tuning was eliminated; (2) diversity in latency tuning was eliminated but diverse spike count tuning remained intact; and (3) diversity in spike count tuning was eliminated but diverse latency tuning remained intact (STAR Methods). These four sets of input are illustrated in Figure 5E (scatter dot size indicates the number of input spikes arriving at a given latency for each stimulus amplitude). In the absence of diverse latency tuning, all EAs fire at approximately the same time across stimulus amplitudes. Similarly, in the absence of diverse spike count tuning, most EAs fire a maximal number of spikes even at relatively low stimulus amplitudes. Consistent with this, removal of diversity in both latency tuning and spike count tuning resulted in postsynaptic responses that were near maximal across a wide range of stimulus amplitudes for both SGCs and DGCs (Figure 5F-H, blue).

The relative importance of diversity in latency tuning versus diversity in spike count tuning depended on the temporal window of postsynaptic integration as set by the timing of inhibition (Figure S4). When inhibition onset was rapid, removing diversity in latency tuning had the major impact on model responses (Figure 5F, purple). In the absence of inhibition, removing diversity in spike count tuning had the major impact on model responses (Figure 5G, orange). For intermediate values of inhibition delay, graded responses could be supported by either diversity in latency tuning or independent spike count tuning (Figure 5H). These results suggest that narrower versus wider temporal integration windows render DGCs and SGCs differentially sensitive to information contained in the relative timing versus the overall number of EA spikes, respectively. Sensitivity of peak response amplitude to the relative timing of EA inputs can be explained by powerful sensory-evoked inhibition and steep rising sensory-evoked excitation (both notable features of DGCs). Assuming that inhibition is triggered by the first EA input to a granular cell (see **Discussion**), a decrease in the relative timing of subsequent EA inputs will result in the postsynaptic response reaching a more depolarized level before being truncated by inhibition (Figure 5I).

### Comparing ELL zones provides additional evidence for the importance of diverse EA tuning

Some degree of heterogeneity in response properties across a neuronal population is likely inevitable, raising the question of whether the diverse EA tuning described here can truly be considered a biological specialization? In addition to the EAs studied here (termed A-type), mormyrid fish also possess additional B-type receptors sensitive to object-induced changes in both the amplitude and shape of the EOD waveform (von der Emde and Bleckmann, 1992, 1997). B-type EAs project to granular cells located within an adjacent sub-region of the ELL known as the dorsolateral zone (DLZ) (Bell et al., 1989). Although circuitry, cell types, and CD input appear generally similar between the MZ and DLZ, prior work has shown that spike threshold is highly uniform across B-type EAs (Bell, 1990b). If diverse tuning is essential for the postsynaptic readout of information conveyed by A-type EAs, the absence of such diversity in B-type EAs would be expected to manifest as differences in postsynaptic responses in the MZ versus the DLZ.

As an initial test of this hypothesis, we compared field potential responses evoked by identical modulations of electrosensory stimulus amplitude for series of closely-spaced electrode penetrations that passed through somatopically aligned regions of the MZ and the DLZ. The early negative component of such field potentials reflects excitation in the granular layer evoked by EAs innervating a small region of the skin (Bell et al., 1992; Gomez et al., 2004; Grant et al., 1998; Sawtell and Williams, 2008). In the MZ, field potential onset latency decreased while the amplitude exhibited prominent grading as a function of stimulus amplitude (Figure 6A). Such amplitude grading can be understood in the same terms as granular cell postsynaptic responses, i.e. as due to an increase in the summed input of a local population of EAs. In the DLZ, field potential onset latency also decreased, however, the amplitude of the field potential exhibited little grading as a function of stimulus amplitude (Figure 6B). To confirm that the difference in response amplitude changes across zones were not due to differences in overall sensitivity to sensory input, we plotted peak field potential amplitude as a function of peak latency across stimulus amplitude (Figure 6C-F). The slope of this relationship was about four times greater in MZ compared to DLZ, though the range of latency shifts was similar (Figure 6F-H). We hypothesize that the lack of amplitude grading in the DLZ is due to the lack of diverse tuning in B-type EAs, similar to the results obtained in the model with simulated EA populations lacking diverse tuning. These results further support the functional importance of diverse tuning in A-type EAs and motivate future studies of the postsynaptic readout of B-type EA input in the granular cells of the DLZ.

**Figure 6.**
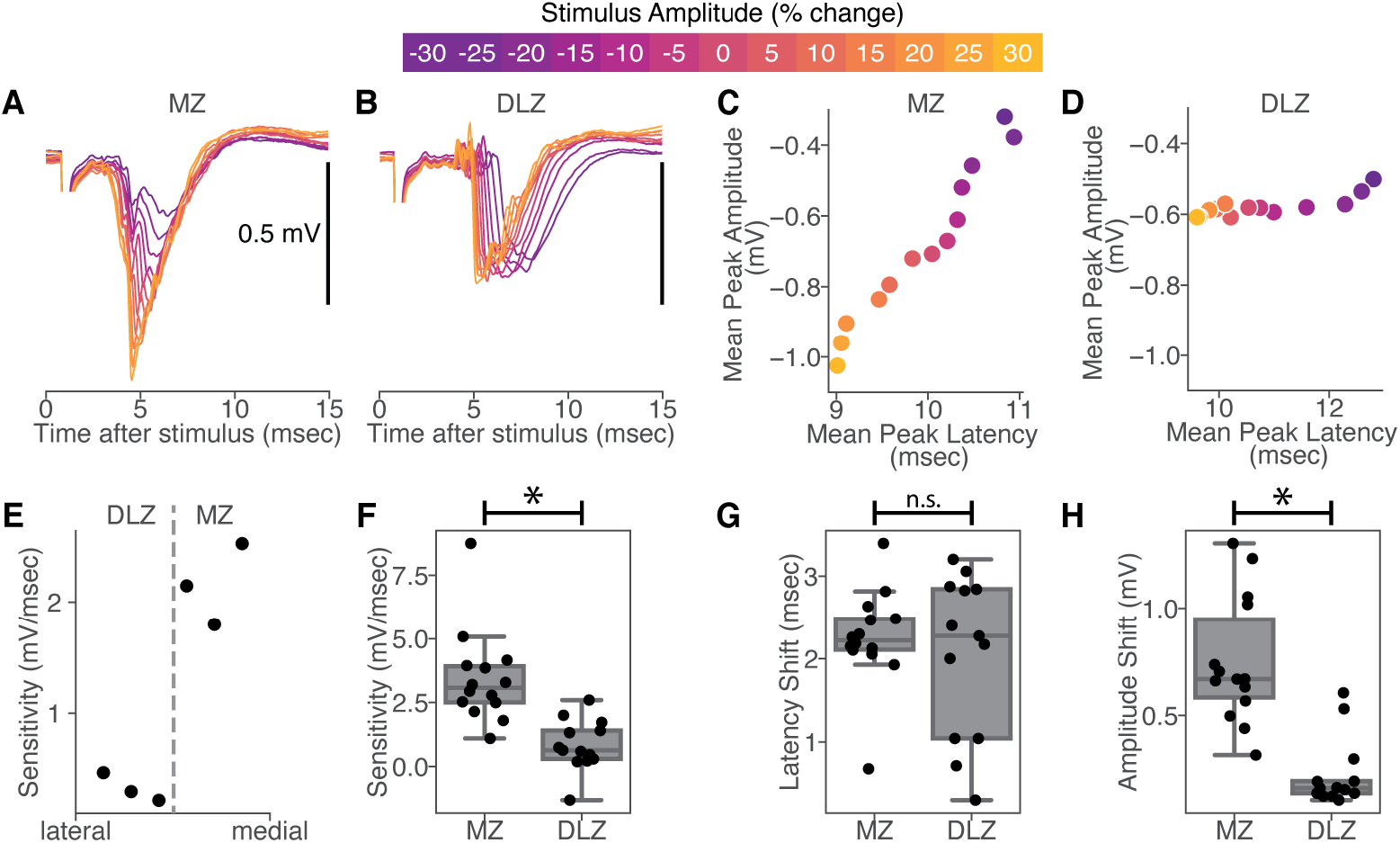
Differences in EA tuning variability result in different stimulus readouts in MZ and DLZ. (A) Local Field Potential (LFP) in the medial zone (MZ) of ELL in response to different stimulus amplitudes (color bar, top). (B) Same as in A, but for the dorsolateral done (DLZ). (C) Scatterplot of LFP amplitude versus LFP peak latency relative to stimulus onset for MZ. (D) Same as in C, but for DLZ. Shifts in stimulus amplitude cause shifts in peak latency for both zones, but peak amplitude changes are greater in the MZ. (E) Sensitivity of LFP amplitude to latency shifts for a series of closely spaced electrode penetrations at different mediolateral locations spanning the MZ-DLZ border, as determined by shifts in receptive location on the skin. Note the abrupt jump in sensitivity around the zonal border. (F) Sensitivity of LFP amplitude to latency shifts is greater in MZ compared to DLZ (t(25)=4.6, p< 0.001; n = 14 MZ sites, n = 13 DLZ sites). Boxplots denote 25%, 50%, and 75% quartiles (with errorbars denoting the rest of the distribution, excluding outliers). (G) LFP peak latency in MZ and DLZ shifts similar amounts across stimulus amplitude (t(25)=0.63, p=0.53; n = 14 MZ sites, n = 13 DLZ sites). (H) LFP amplitude changes are greater in MZ versus DLZ (t(25)=5.8, p< 0.001; n = 14 MZ sites, n = 13 DLZ sites).

## DISCUSSION

Compared to the many studies characterizing the encoding of sensory information by spike latency shifts, there have been relatively few tests of whether and how such information is actually utilized by postsynaptic neurons. The active electrosensory system is advantageous in this regard because latency decoding is hypothesized to occur in granular cells located just one synapse from the sensory periphery. In addition, central reference signals hypothesized to perform the decoding can be easily monitored and manipulated *in vivo*. However, until now, their small size and dense packing have precluded *in vivo* recordings from individual granular cells. Using *in vivo* whole-cell recordings we demonstrate that granular cells exhibit large stimulus-evoked changes in postsynaptic response amplitude, consistent with prior hypotheses that latency shifts in EAs are transformed into changes in response amplitude (Bell, 1990a). Although granular cells integrate EA input with motor CD signals providing a precise reference signal related to stimulus onset, such signals appear to gate or enhance responses rather than being strictly required to decode sensory input. Instead, we find that diverse latency and spike count tuning across the EA population gives rise to graded information about stimulus amplitude in the form of a rate code. Modeling indicates that sensory-evoked postsynaptic responses in two distinct subclasses of granular cells can be explained, without a fixed reference signal, simply by summation of excitatory input from a small number of converging EA inputs. Finally, a brief integration window set by powerful sensory-evoked inhibition renders DGCs highly sensitive to information contained in the relative latency of EA spikes, while the absence of strong inhibition renders SGCs more sensitive to total spike count.

### Comparison to latency readout schemes proposed for other systems

One common proposal for reading out latency codes relies on fixed reference signals related to stimulus onset. CD signals are hypothesized to provide a reference signal for reading out latency codes in systems where the timing of sensory input is determined by the behavior of the animal (Crapse and Sommer, 2008; Cury and Uchida, 2010; Hall et al., 1995; Smear et al., 2011), for example saccades in vision, sniffing in olfaction, active touch in somatosensation, or the EOD in electrolocation. Alternatively, fixed reference signals could be contained in the sensory input itself, for example by subsets of sensory neurons with short, stereotyped response latencies (Brasselet et al., 2012; Chase and Young, 2007; Gollisch and Meister, 2008). Other readout schemes rely on postsynaptic specializations such as specialized learning rules for tuning synaptic strength (Gutig and Sompolinsky, 2006; Thorpe et al., 2001), or competitive interactions mediated by recurrent inhibition (Haddad et al., 2013; Stern et al., 2018; Zohar and Shamir, 2016). The present findings are notable in that no fixed reference signal is required. Furthermore, diverse tuning across sensory input populations has been reported in numerous systems, see e.g. (Bale et al., 2013; Goldberg, 2000; Raman et al., 2010), suggesting that the mechanism described here may be of general relevance for understanding how latency codes are utilized by the brain. In the mammalian auditory system, for example, there is wide variation in the sensitivity of auditory nerve fibers that co-varies with spontaneous firing rate, axonal morphology, and transcriptional patterns, and has been functionally linked to the wide dynamic range of human hearing (Liberman, 1982; Liberman and Oliver, 1984; Petitpre et al., 2018; Viemeister, 1983; Winter et al., 1990).

Although the origin of the diverse tuning observed across A-type electroreceptors is not known, a prior electrophysiological study noted that A-type electroreceptors exhibit widely varying spike thresholds and an electron microscopy study observed notable variation in the area of the outer membrane of A cells (Bell, 1990b; Bell et al., 1989). Differences in types, distributions, or densities of voltage-gated channels across A-type electroreceptor cells or their afferent fibers have not been investigated but could also contribute to diverse responses. Another question for future studies is whether convergence of A-type EAs onto granular cells is random (as in our model) or subject to some forms of tuning or optimization. Although we did not systematically examine non-random connectivity, certain rules might be expected to enhance postsynaptic responses. For example, convergence of EAs with intersecting latency tuning curves is expected to give rise to non-monotonic postsynaptic responses (i.e. maximal responses at the intersection point). Response grading could potentially be enhanced if granular cells avoided pooling such EAs.

### Limitations and functional implications

An important limitation of the present study was our inability to reliably measure the spiking output of granular cells. Spike threshold, as well as additional postsynaptic non-linearities, could enhance sensitivity to small changes in EOD amplitude. Nevertheless, the observation that subthreshold responses to electrosensory stimuli in E-type output cells were similar to those in granular cells, support our conclusions that CD enhances, but is not required for, encoding electrosensory stimulus amplitude (Figure S3). A second limitation relates to the use of spatially uniform electrosensory stimulation. These simple stimuli greatly facilitated quantitative comparisons, which would have been difficult with local stimuli given the difficulty of maintaining stable recordings from small granular cells while mapping receptive fields. However, they obviously restricted us from studying potentially important effects of the spatial structure of electrosensory input. Due to the inherent blur in electrical images, EAs converging onto a given granular cell probably convey similar signals. However, LMIs likely receive electrosensory input from large regions of the body surface and may perform spatial computations such as lateral inhibition (Han et al., 2000; Meek et al., 2001). Given the apparent differences in sensory-evoked inhibition between DGCs and SGCs, these two sub-classes may differ not only in the temporal profiles of their sensory responses but also in their spatial tuning. Additional key questions relate to synaptic and functional connectivity patterns linking granular cells to the rest of the ELL. While both DGCs and SGCs send axons to the superficial layers of the ELL where they contact several anatomically and functionally distinct neuronal sub-classes, the details of their connectivity patterns are not known (Hollmann et al., 2016; Meek et al., 1999).

The observation that precise central reference signals are present in granular cells raises the question of what advantages, if any, are conferred by the readout scheme proposed here. In contrast to the fixed temporal window provided by corollary discharge, sensory-evoked inhibition in DGCs provides a window for reading out relative latency information that shifts along with the sensory input. This may provide a means for maintaining sensitivity to small latency shifts, such as those due to prey (Gottwald et al., 2018), superimposed on larger shifts due to the animals’ own behavior or large environmental features (Chen et al., 2005; Sawtell and Williams, 2008; Sawtell et al., 2006). Anatomical studies have shown that the myelinated dendrites of LMI cells form large GABAergic synapses onto granular cells. LMI dendrites lack conventional synaptic inputs and are hypothesized to be activated directly by depolarization within granular cells via ephaptic coupling (Han et al., 2000; Meek et al., 2001). This unusual synaptic organization appears well-suited to mediate the rapid and powerful inhibition observed in DGCs. Whereas DGCs, LMI cells, and latency coding in EAs appear to be specializations of the active electrosensory system (all three are absent from an adjacent zone of the ELL that subserves the more evolutionarily ancient passive electrosense), SGCs are found in both systems (Bell et al., 2005). Based on our finding that SGCs are mainly sensitive to spike count while DGCs are sensitive to input timing, it is tempting to speculate that DGCs and LMIs are adaptations for utilizing temporal information associated with the evolution of the active electrosense. Finally, encoding stimulus amplitude independent of CD could enable a dual role for the active electrosensory system in utilizing information derived from the EODs of other fish. Consistent with such a function, one prominent close-range electrocommunication behavior, the echo response, appears to be mediated by the active electrosensory system rather than by separate electrocommunication pathways (Russell et al., 1974).

## Acknowledgements

This work was supported by grants from the NIH (NS075023), NSF (IOS-1656354) and Irma T. Hirschl Trust to N.B.S. and a Simons Society of Fellows Junior Fellowship to K.P. We thank C. Bell for helpful discussions, L.F. Abbott for discussions and comments on the manuscript, and Federico Pedraja for assistance with Figure 1.

## Author Contributions

K.P. and N.B.S. conceived of the project and designed the experiments. K.P. performed the experiments, analyzed the data, and performed the modeling. K.P. and N.B.S. wrote the manuscript.

## Supplemental Figures

**Figure S1.**
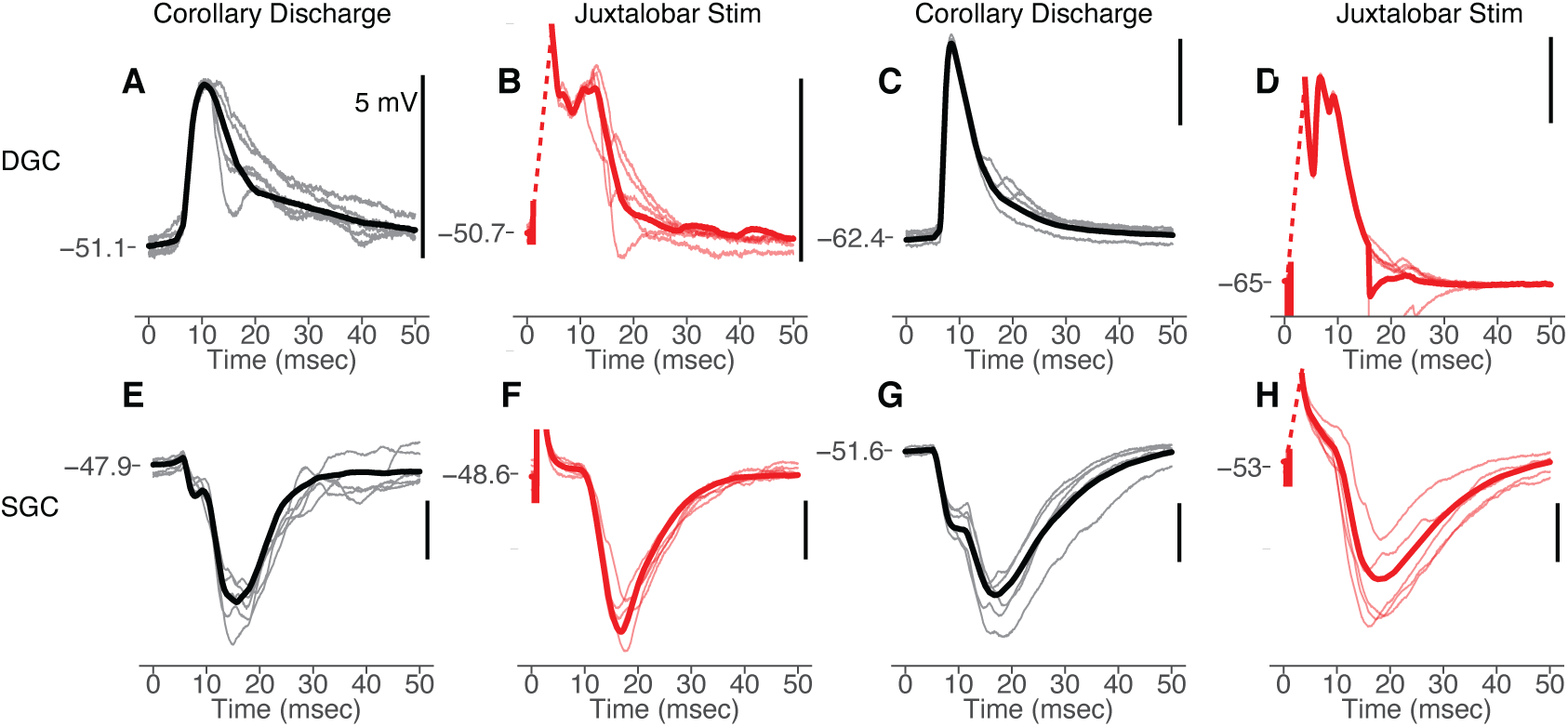
Electrical microstimulation of the medial juxtalobar nucleus evokes responses in DGCs and SGCs that mimic electric organ corollary discharge responses. Related to Figure 2. (A) Subthreshold response of an example DGC time-locked to the command to discharge the electric organ (t=0). Gray: five overlaid trials; Black; trial average. Since paralysis blocks emission of the EOD, responses time-locked to the EOD motor command are due to electric organ corollary discharge input. (B) Response of the same DGC time-locked to the electrical stimulation of the medial juxtalobar nucleus (t=0). Pink: five overlaid trials; Red; trial average. Juxtalobar stimulation evokes an EPSP at a short latency, similar to the corollary discharge response in this cell. (C, D) Same display as in (A) and (B) for a second example DGC. (E) Membrane potential response of an example SGC time-locked to the command to discharge the electric organ (t=0). Gray: five overlaid trials; Black; trial average. (F) Response of the same SGC time-locked to the electrical stimulation of the medial juxtalobar nucleus (t=0). Pink: five overlaid trials; Red; trial average. Juxtalobar stimulation evokes an IPSP, similar to the corollary discharge response in this cell. Note, that the inflection on the falling phase of the IPSP, presumably reflecting corollary discharge driven excitation (see **Figure 2F**), is not reproduced by juxtalobar stimulation, suggesting the possibility of an additional source of corollary discharge excitation to SGCs. (G, H) Same display as in (E) and (F) for a second example SGC.

**Figure S2.**
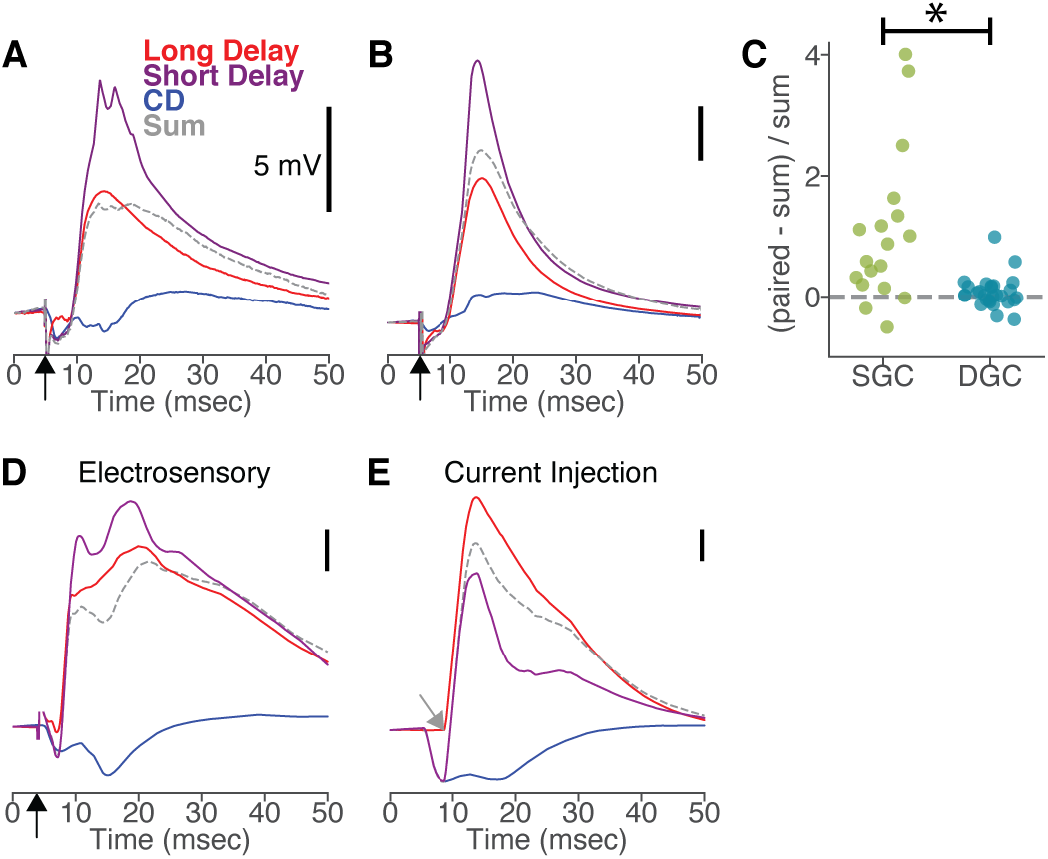
Supralinear summation of electrosensory and corollary discharge responses in SGCs. Related to Figure 3. (A) Average responses of an example SGC to the electrosensory stimulus (black arrow) at baseline amplitude delivered either at a long delay (red) or short delay (purple) relative to the EOD motor command. Blue: average response to corollary discharge (CD) input alone (time-locked to the EOD motor command). The predicted response (sum; gray dashed) was calculated by adding the electrosensory response at a long delay to the CD response. The response to an electrosensory stimulus at a short delay relative to the EOD is much greater than the predicted linear sum of the EA and CD inputs alone. (B) Same as in (A) but for a different example SGC showing another case of supralinear summation between electrosensory and CD input. (C) The difference between the predicted and the short delay response peak amplitudes (normalized by the predicted amplitude) was significantly larger for SGCs (green; 0.7 ± 1.9 n=19) than for DGCs (blue; 0.07 ± 0.26, n = 28; t(45)= 4.02, p< 0.001 with an SGC outlier at −5.3 omitted). (D) Same display as in (A) but for a different SGC. (E) Response of the same SGC as in (D) but substituting an intracellularly injected current waveform (gray arrow) for the electrosensory stimulus. Supralinear summation was not observed between the intracellularly-evoked depolarization and the corollary discharge. Similar results were obtained in a total of 5 SGCs, suggesting that the supralinear summation is not primarily due to intrinsic effects, such as active conductances in SGCs, or that such effects are not engaged by somatic current injections.

**Figure S3.**
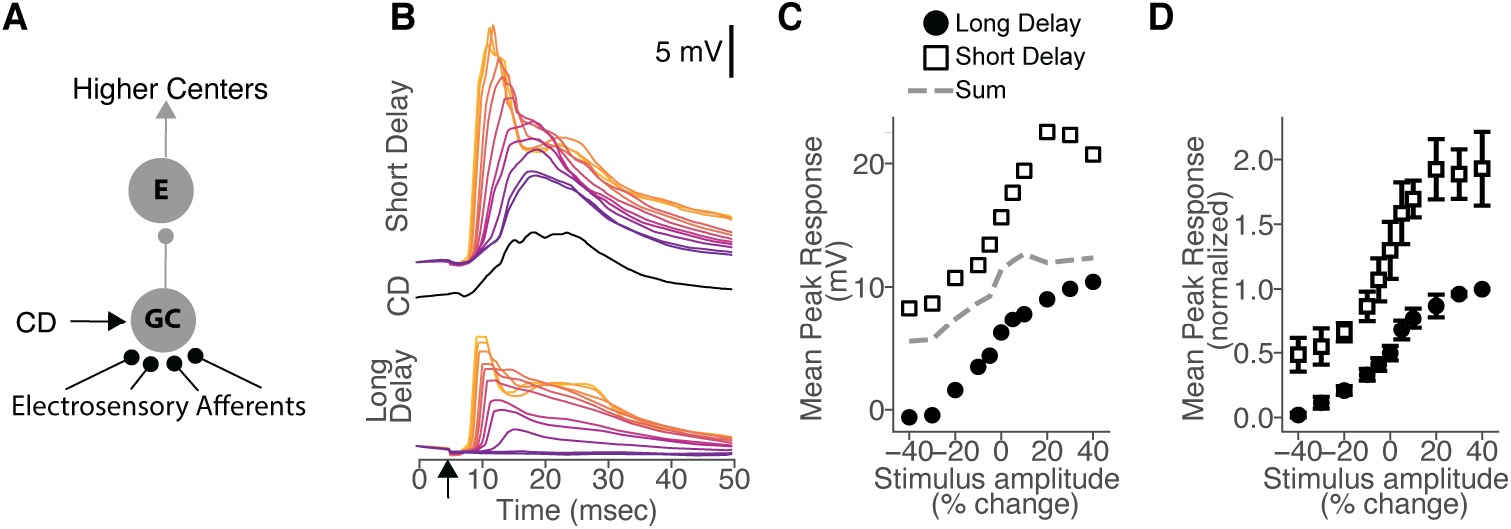
Supralinear summation of electrosensory and corollary discharge responses in GC output. Related to Figure 3. (A) Granular cells (GC) integrate convergent input from peripheral electroreceptor afferents with a central corollary discharge (CD). GCs relay electrosensory input to several classes of neurons located in the superficial layers of the ELL, including the E-type output cells (E) recorded here. E-type output cells also receive separate sources of CD input not shown. (B) Responses of an example E-type output cell across amplitudes to stimuli delivered either at a short (top) or long (bottom) delay relative to the EOD motor command (each waveform is an average across trials). Middle: average response to CD input alone. (C) Mean peak response versus stimulus amplitude for the cell shown in (D) in long delay (filled circle) versus short delay (open square) conditions. The predicted response at a short delay (‘sum’, gray dashed) was calculated by adding the long delay response at each stimulus amplitude to the CD response. (D) Average peak response amplitude versus stimulus amplitude under long delay (filled circle) versus short delay (open square) conditions across E-type output cells (n = 5; mean ± SEM). Responses within each cell were normalized by the maximum long delay response amplitude before averaging across cells. The delay between the electrosensory stimulus and the command had a significant effect on peak response amplitude, F(1,88) = 160, *P* < 0.001 (two-factor repeated measures ANOVA). The population mean response changed, across minimum to maximum stimulus amplitude, by 9.2 ± 4.0 mV at a long delay and 12.5 ± 4.8 mV at a short delay. Out of 4 output cells with at least 5 trials in every condition, 4 had a significant effect of stimulus delay on raw peak response amplitude (p< 0.001). Significant effects of stimulus amplitude and delay, but not delay-by-amplitude interaction at P < 0.001 on the mean peak response amplitudes across cells.

**Figure S4.**
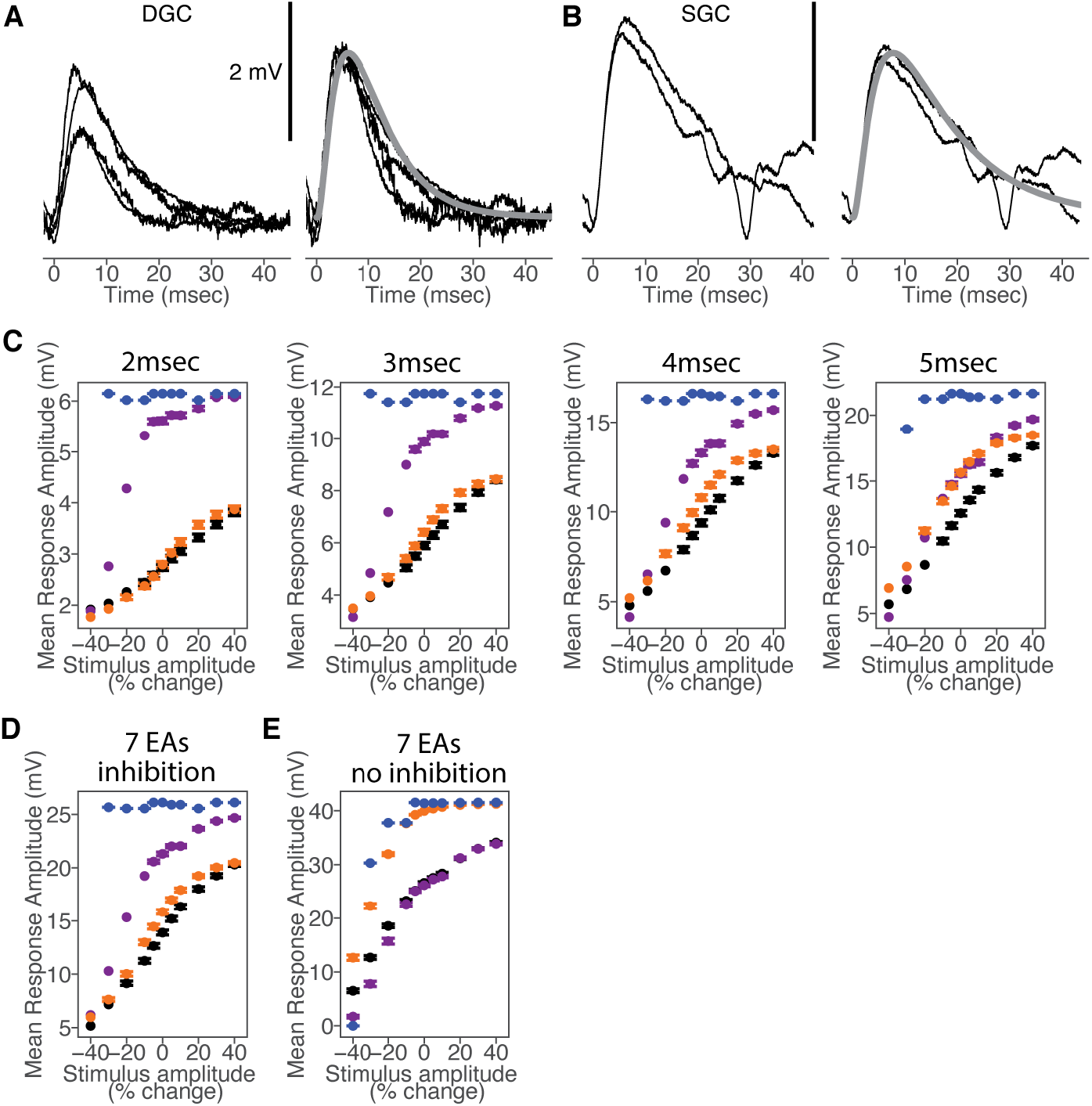
Effects of varying model parameters on granular cell postsynaptic responses. Related to Figure 5. (A) Left: Average responses evoked by a near-threshold electrosensory stimulus in DGCs (each trace is the average response from a different DGC). These data were used to fit synaptic time constants of afferent input in the model. Right: The same EPSPs as shown to the left, but normalized to the peak amplitude and overlaid with the normalized model EPSP for DGCs (gray; 3 mV peak amplitude) to compare the temporal dynamics. DGC EPSP kinetics were a bit faster than for SGCs. To fit this difference, model DGCs differed from model SGCs, but only in the membrane capacitance (6 pF for DGCs and 12 pF for SGCs). (B) Same as in (A) but for 2 SGCs (left) and the normalized SGC model EPSP overlaid (right; gray). (C) Impact of diverse latency versus spike count tuning in EAs on model granular cell responses depends on inhibition delay. Peak response amplitude versus stimulus amplitude across model GCs (12 pF membrane capacitance; n = 200 different EA input populations; mean ± SEM) with 4 EA inputs under each condition shown in **Figure 5E**(Black, subsampled recorded EA data; Orange, diverse latency tuning is preserved while diverse spike count tuning is removed; Purple, diverse latency tuning is removed while diverse spike count tuning is preserved; Blue, diverse latency and spike count tuning are both removed from the input population). Removing diversity in latency tuning from the input population has the largest impact when inhibition is minimally delayed (2 msec, left). (D) Same as in (C), but for model GCs (n = 200 different EA input populations; mean ± SEM) with 7 EA inputs and inhibition delayed by 4 msec relative to the first EA spike. Increasing input number increases the overall magnitude of postsynaptic responses, but does not qualitatively impact the effects of manipulating diversity in EA tuning (compare to results for 4 EA inputs shown in **Figure 5H**). (E) Same as in (D), but for model GCs (n = 200 different EA input populations; mean ± SEM) with no inhibition. Increasing input number increases the overall magnitude of postsynaptic responses, but again does not qualitatively impact the effects of manipulating diversity in EA tuning (compare to results for 4 EA inputs shown in **Figure 5G**).

## STAR Methods

### Contact for Reagent and Resource Sharing

Further information and requests for resources and reagents should be directed to and will be fulfilled by the Lead Contact Nathaniel Sawtell

### Experimental Model and Subject Details

Male and female Mormyrid fish (7-12 cm in length) of the species *Gnathonemus petersii* were used in these experiments. Fish were housed in 60 gallon tanks in groups of 5-20. Water conductivity was maintained between 70-150 microsiemens both in the fish’s home tanks and during experiments. All experiments performed in this study adhere to the American Physiological Society’s Guiding Principles in the Care and Use of Animals and were approved by the Institutional Animal Care and Use Committee of Columbia University.

For surgery to expose the brain for recording, fish were anesthetized (MS:222, 1:25,000) and held against a foam pad. Skin on the dorsal surface of the head was removed and a long-lasting local anesthetic (0.75% Bupivacaine) was applied to the wound margins. A plastic rod was cemented to the anterior portion of the skull to secure the head. The posterior portion of the skull overlying the ELL was removed. The valvula cerebelli was reflected laterally to expose the eminentia granularis posterior and the molecular layer of the ELL, facilitating whole-cell recordings. Gallamine triethiodide (Flaxedil) was given at the end of the surgery (∼20 μg/cm of body length) and the anesthetic was removed. Aerated water was passed over the fish’s gills for respiration. Paralysis blocks the effect of electromotoneurons on the electric organ, preventing the EOD, but the motor command signal that would normally elicit an EOD continues to be emitted at an average rate of 2 to 5 Hz.

### Electrophysiology

The EOD motor command signal was recorded with a Ag-AgCl electrode placed over the electric organ. The command signal is the synchronized volley of electromotoneurons that would normally elicit an EOD in the absence of neuromuscular blockade. The command signal lasts about 3 ms and consists of a small negative wave followed by three larger biphasic waves. Onset of EOD command was defined as the negative peak of the first large biphasic wave in the command signal.

For *in vivo* whole-cell recordings, electrodes (8-15 MΩ) were filled with an internal solution containing, in mM: K-gluconate (122); KCl (7); HEPES (10); Na2GTP (0.4); MgATP (4); EGTA (0.5), and 0.5%–1% biocytin (pH 7.2-7.4, 280-315 mOsm). No correction was made for liquid junction potentials. Only cells with stable membrane potentials more hyperpolarized than −45 mV were analyzed. Membrane potentials were recorded and filtered at 3-10 kHz (Axoclamp 2B amplifier, Axon Instruments) and digitized at 20-40 kHz (CED micro1401 hardware and Spike2 software; Cambridge Electronics Design, Cambridge, UK).

### Electrosensory stimulation

The EOD mimic was a 0.2 ms duration square pulse delivered between an electrode in the stomach and another positioned near the electric organ in the tail. In between recordings, the EOD mimic was presented at a baseline amplitude of 350 μA (stomach electrode negative) at the output of the stimulus isolation unit. To characterize neural tuning curves, stimulus amplitude was varied from +40% to – 40% from the baseline amplitude. To characterize interactions between electrosensory and corollary discharge inputs, responses were compared across conditions in which the EOD mimic was presented at a delay of either 50 ms (long delay) or 4.5 ms (short delay) following the EOD command.

### Juxtalobar stimulation

The precise location of the juxtalobar nucleus was determined by monitoring corollary discharge-evoked field potentials with extracellular recording electrodes. Low resistance, broken-tip glass capillary microelectrodes filled with 3M NaCl were used for recording field potentials and for electrical stimulation of the juxtalobar nucleus (stimulus duration was a single square wave pulse with a duration of 200 microseconds). The electrodes were directed at angles of about 45 degrees with respect to the mid-sagittal plane and with a slight posterior to anterior direction. With the valvula reflected, entry points for the electrode tracks were just dorsal to the anterior tip of the exposed electrosensory lobe molecular layer. The corollary discharge-driven field potential characteristic of the juxtalobar nucleus was recorded in such tracks at depths of 800 to 1500 microns below the surface (depending on fish size and exact tilt). Prior to searching for and recording granular cells, juxtalobar nucleus electrode placement was confirmed by the ability of a near-threshold stimulus to evoke characteristic field potential responses in the granular layers of ELL (Mohr et al., 2003). Minimum stimulus thresholds for evoking responses in ELL were 2-4 uA. For evoking responses in granular cells, a near threshold stimulus of 5uA was used.

### Histology and morphological reconstructions

After recording, fish were deeply anesthetized with a concentrated solution of MS:222 (1:10,000) and perfused through the heart with a teleost Ringer solution followed by a fixative, consisting of 4% paraformaldehyde in 0.1 M phosphate buffer. The brains were postfixed for 12-24 hours, cryoprotected with 30% sucrose, and sectioned at 60 μm on a cryostat. Sections were subsequently processed with an Alexa Fluor 488 Streptavidin complex (Jackson Immuno Research Laboratories; Antibody ID - AB_2337249; at 1:500) to label the biocytin filled cells, and DAPI (Sigma Aldrich # D9542; at 1:1000) and NeuroTrace 640 (ThermoFisher Scientific # N21483; at 1:500) to visualize the layers of ELL. Sections were then mounted on slides, dried of excess PBS, and coverslipped with either VectaShield Antifade (Vector Laboratories # H-1000-10) or Molecular Probes™ ProLong™ Diamond Antifade Mountant (Fisher Scientific ; Molecular Probes™ P36965). Morphologically recovered neurons were inspected and subsequently photographed using a confocal microscope (Inverted Nikon A1R point-scanning laser confocal microscope with high-sensitivity GaAsP detectors) with either a 20x air objective or a 40x oil immersion objective. Images were collapsed along the Z-dimension implementing the maximum brightness per pixel. Each fluorescence channel was pseudo-colored as specified in Figure 2.

### Modeling

#### Circuit Architecture

The circuit consisted of one model granular cell receiving input from a small number (4 used in the main results, 4-7 tested in supplemental) simulated afferents and one simulated inhibitory input.

#### Model granular cell

Each model granular cell is described by a single compartment with a membrane potential that evolves according to:

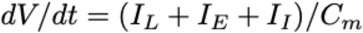

Membrane capacitance (C_M_) values were chosen such that the model EPSP evoked by a single presynaptic input matched near threshold electrosensory responses of DGCs and SGCs (Figure S4). These values were 6 pF for DGCs and 12 pF for SGCs. The leak current (I_L_) is described by:

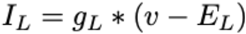

where v is the membrane potential of the model granular cell, the leak conductance g_L_ = 1nS and the reversal potential E_L_ = −70mV. These values were consistent with a prior *in vitro* study of ELL granular cells (Zhang et al., 2007).

#### Model synapse

Each synaptic current (I_E_ and I_I_) is described by the standard equation:

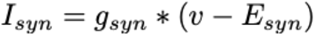

where the reversal potential of the excitatory conductance is 0mV and the reversal potential of the inhibitory conductance is −90 mV. The timecourse of the excitatory conductance (g_E_) follows a double exponential with a rise time constant τ_E1_ = 4 ms and a decay time constant τ_E2_ = 1 ms. The conductance parameter ‘s’ is increased by w_e_ upon a spike event in a presynaptic electrosensory afferent and decays exponentially with time constant τE2. The excitatory conductance then evolves according to:

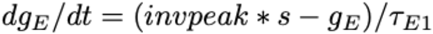

where

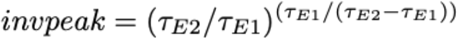

The inhibitory conductance (g_I_) is increased by wI upon a spike event in the stimulated presynaptic LMI and decays exponentially with time constant τ_I_ = 10ms.

#### Electrosensory afferent (EA) input

For each model granular cell, a set of 4 EA inputs were either subsampled from recorded afferent data or simulated. The main results were not affected by the exact number of EA inputs to each model granular cell (Figure S4), for values within a plausible range based on past studies of the ELL (Bell, 1990a; Bell et al., 2005; Zhang et al., 2007). We simulated afferents by first calculating an exponential tuning curve of first spike latency versus stimulus amplitude. We randomly selected each set of exponential fit parameters from a multivariate Gaussian distribution that was fit to the recorded afferent data (Figure 4E-G). We then added multiple spikes according to the mean spike offsets calculated in recorded EAs (Figure 4I). We constructed three different EA input conditions to separately test the effects of diverse latency tuning and diverse spike tuning and the combined effect of both (Figure 5E). To test the effect of diverse latency tuning on model granular cell responses (purple in Figure 5E-H), we constructed a model EA tuning curve using the afferent population mean value for each latency tuning parameter, which was held constant, and added subsequent spikes with an offset. We then randomly selected spike count tuning from recorded EAs to mask spikes in the latency tuning curve. To test the effect of diverse spike count tuning on granular cell responses (orange in Figure 5E-H), we constructed model EA tuning curves by randomly selecting each set of exponential fit parameters from a multivariate Gaussian distribution that was fit to the recorded afferent data. We then added subsequent spikes with an offset up to a maximum spike latency calculated from the recorded EA data (11 ms). To test both diverse latency and spike count tuning (blue in Figure 5E-H), we constructed a model EA tuning curve using the afferent population mean value for each latency tuning parameter, which was held constant, and added subsequent spikes with an offset up to a maximum spike latency calculated from the recorded EA data (11 ms).

#### LMI inhibitory input

We simulated electrosensory-evoked inhibitory input to the model by specifying an input spike time with a constant onset latency relative to the earliest simulated EA input spike (we tested a range of onset latency from 2-5 msec; Figure S4). The weight of the inhibitory synapse (wI) was varied as a sigmoid function of the membrane potential of the post-synaptic granular cell (v) at the time of the LMI spike, which provided a nice fit to the data (Figure 5C,D). This is meant to approximate the proposed ephaptic mechanism of LMI recruitment suggested by prior work (Han et al., 2000; Meek et al., 2001). However, qualitatively similar results were obtained if the strength of inhibitory input was held constant as long as it was strong enough to truncate the response and suppress its normal peak (data not shown).

#### Model Simulation

All simulations were done in Python3 with the BRIAN 2 simulator package (Stimberg et al., 2019). The model had a time step of 0.1 ms and was simulated for 50 ms in response to each electrosensory stimulus. Model GC responses were then quantified by measuring the peak membrane potential as was done for the real GCs recorded in this study. Each model cell was a simulation with a different EA input population.

### Software used and general statistical methods

Data were analyzed offline using Python 3 software. Biophysical models were simulated using the BRIAN 2 simulator package (Stimberg et al., 2019) (as noted above) and analyzed using Python 3. No statistical methods were used to predetermine sample size. The experimenters were not blinded to the condition during data collection or analysis. Statistical test identity is indicated along with each result. Differences were considered significant at p < 0.05.

### Data and Code Availability

Data will be available via GNode. Data analysis code will be available at github.

